# First draft genome assembly of an iconic clownfish species (*Amphiprion frenatus*)

**DOI:** 10.1101/205443

**Authors:** Anna Marcionetti, Victor Rossier, Joris A. M. Bertrand, Glenn Litsios, Nicolas Salamin

## Abstract

Clownfishes (or anemonefishes) form an iconic group of coral reef fishes, particularly known for their mutualistic interaction with sea anemones. They are characterized by particular life history traits, such as a complex social structure and mating system involving sequential hermaphroditism, coupled with an exceptionally long lifespan. Additionally, clownfishes are considered to be one of the rare group to have experienced an adaptive radiation in the marine environment.

Here, we assembled and annotated the first genome of a clownfish species, the tomato clownfish *(Amphiprion frenatus*). We obtained a total of 17,801 assembled scaffolds, containing a total of 26,917 genes. The completeness of the assembly and annotation was satisfying, with 96.5% of the *Actinopterygii* BUSCOs (<u>B</u>enchmarking <u>U</u>niversal <u>S</u>ingle-<u>C</u>opy <u>O</u>rthologs) being retrieved in *A. frenatus* assembly. The quality of the resulting assembly is comparable to other bony fish assemblies.

This resource is valuable for the advancing of studies of the particular life-history traits of clownfishes, as well as being useful for population genetic studies and the development of new phylogenetic markers. It will also open the way to comparative genomics. Indeed, future genomic comparison among closely related fishes may provide means to identify genes related to the unique adaptations to different sea anemone hosts, as well as better characterize the genomic signatures of an adaptive radiation.

## Introduction

Clownfishes (or anemonefishes; subfamily Amphiprioninae, genera *Amphiprion* and *Premnas*) are an iconic and highly diverse group of coral reef fishes. They are part of the damselfish family (Pomacentridae) and they include 28 described species [Ollerton *et al.* 2007]. Their distribution spans the whole tropical belt of the Indo-West Pacific Ocean, but their highest species richness is situated in the Coral Triangle region, where up to nine clownfish species have been observed in sympatry [Elliott and Mariscal 2001].

One distinctive characteristic of this group is the mutualistic interaction they maintain with sea anemones [Fautin and Allen 1997]. While all species of the clade are associated with sea anemones, there is a large variability in host usage within the group. Indeed, some species are strictly specialist and can interact with a unique species of sea anemones, while others are generalists and can cooperate with a large number of hosts [Ollerton *et al.* 2007]. Studies have been conducted to understand both the process of host selection used by clownfishes [e.g. Arvedlund *et al.* 1999, Elliott *et al.* 1995, Elliott and Mariscal 2001, Huebner *et al.* 2012] and the mechanisms granting them protection from sea anemones toxins [reviewed in Mebs 2009]. However, we do not have yet a full answer for these questions. In particular, the genomic bases of these mechanisms remain poorly understood.

Clownfishes also have particular life-history traits and strategies compared to other damselfishes and most other coral reef fishes. They display an outstanding lifespan, with around 30 years estimated for *A. percula*. This lifespan is twice as long as any other damselfishes and six times greater than the expected longevity for a fish of their size [Buston and Garcia, 2007]. Moreover, clownfishes live in complex social structures within the anemones and are protandrous hermaphrodites [Buston 2003]. Studies have been conducted to understand the maintenance of this social structure [e.g. Buston 2004, Mitchell 2003, Hattori 2000] and the mechanisms involved in sex change [e.g. Kim *et al*. 2010, 2012, 2013, Miura *et al. 2013.* Casas *et al*. 2016].

Litsios *et al.* [2012] proposed that the obligate mutualistic interaction of clownfishes with sea anemones acted as the main key innovation that triggered the adaptive radiation of the group. It was further shown that geographic isolation associated with a rather small dispersal capacity and hybridization played a role in driving the burst of diversification and the adaptive process of this group [Litsios *et al.* 2014, Litsios and Salamin, 2014]. The clownfishes could represent a potentially new and interesting model system for the study of adaptive radiations and could be used to validate the theoretical findings on the dynamics of this process [Gavrilets and Vose 2005, Gavrilets and Losos 2009].

Despite the many and different aspects of clownfishes that are being studied in different fields, the knowledge on their long-term evolution and its underlying genetic bases remains scarce. Yet, advances in next-generation sequencing technologies allow now to obtain genomic information also for non-model organisms. More precisely, the widely used Illumina short reads can be complemented with Pacific Biosciences (PacBio) long reads for hybrid assemblies [Koren *et al.*, 2011, Deshpande *et al.* 2013, Miller *et al.* 2017]. This dual strategy is potentially fruitful as it allows to overcome the errors due to both the repeated regions of the genome that cannot be unambiguously assembled with short reads and the relatively higher error rate with long reads. Indeed, Illumina technology tends to be particularly sensitive to the first kind of error whereas PacBio technology is expected to be more affected by the former one. Additionally, the sequencing of RNA can be used to improve the gene annotation in newly assembled genomes [Denton *et al.* 2014].

In this study, we aimed at obtaining the first draft genome of a clownfish species: the tomato clownfish (*Amphiprion frenatus*). This resource will provide new tools for future investigation of clownfishes particular life-history traits and the study of their mutualism with sea anemones. Additionally, new markers for phylogenetic and population genetics studies can be developed thanks to this draft reference genome. This resource also opens the way to comparative genomics among closely related fishes to identify genes related to the unique adaptations of clownfishes to their different sea anemone hosts. This genomic resource will provide the possibility to link these different fields of research and make a step forward in the understanding of clownfishes ecology and evolution.

## Material and methods

### DNA extraction, library construction and Illumina sequencing

One individual of *A. frenatus* was obtained from a local aquarium shop. Genomic DNA (gDNA) was extracted from about 50 mg of fin tissue using DNeasy Blood & Tissue Kit (Qiagen, Hilden, Germany) following manufacturer’s instructions. The total amount of gDNA was measured using Qubit dsDNA HS Assay Kit (Invitrogen, Thermo Fisher Scientific, Waltham, USA). The integrity of the gDNA was verified with Fragment Analyzer Automated CE System (Advanced Analytical Technologies, Fiorenzuola d’Arda, Italy). A total of 100 ng and 4 μg of gDNA were used for paired-end (PE) and mate pair (MP) library preparation, respectively.

Short-insert (350 bp) PE library was prepared at the Lausanne Genomic Technologies Facility (LGTF, Switzerland) using TruSeq Nano DNA LT Library Preparation Kit (Illumina). Long-insert (3 kb) MP library was prepared at Fasteris SA (Geneva, Switzerland) using the Nextera Mate Pair Library Preparation Kit from Illumina. The concentration, purity and size of the libraries obtained were verified using Agilent Bioanalyzer 2100 (Agilent Technologies, Santa Clara, CA). The PE library was sequenced on 2 lanes of Illumina HiSeq2000 at the LGTF (run type: paired-end reads, read length of 100). The MP library was sequenced on half lane of Illumina HiSeq2500 at Fasteris (run type: paired-end reads, read length of 125 bp).

### DNA extraction, library construction and Pacific Biosciences (PacBio) sequencing

A second individual of *A. frenatus* was obtained from a local aquarium shop. High-molecular-weight gDNA was extracted from about 100 mg of muscle tissue using QIAGEN Genomic-tip 100/G (Qiagen, Hilden, Germany) following manufacturer’s instructions. The total amount of gDNA was measured using Qubit dsDNA HS Assay Kit and the integrity of the gDNA was verified with Fragment Analyzer Automated CE System. The construction of the SMRTbell sequencing libraries and the sequencing were performed at the LGTF from a starting material of 10 μg of gDNA. The SMRTbell libraries were sequenced on eight SMRT cells (Pacific Bioscience) using C2 chemistry on the PacBio RS (Pacific Biosciences) sequencing platform.

### Preprocessing of sequenced reads

Reads quality has a major impact on the quality of the resulting assembly and the use of error-corrected reads increases dramatically the size of the contigs [Salzberg *et al.* 2012]. Two different PE reads correction strategies were therefore performed. The first consisted in correcting raw reads, without prior processing, with ALLPATHS-LG module for the correction of fragment read error (default parameters) [Gnerre *et al.* 2011].

The second strategy consisted of three steps. We removed PE reads that failed the chastity filtering of the Casava pipeline with casava_filter_se.pl (v0.1-1, from http://brianknaus.com/software/srtoolbox/). Remaining PE reads were trimmed using Sickle (v1.29) [Joshi and Fass, 2011], with the following parameters: --qual-threshold 30 --length-threshold 80. Substitutions due to sequencing errors in the trimmed PE reads were corrected with Quake [Kelley *et al*. 2010]. The *k*-mers frequency needed by Quake were obtained with Jellyfish [Marçais and Kingsford, 2011]. A *k*-mer size of *k*=18 was selected according to Quake documentation, which suggests the use of *k*=log(200*GenomeSize)/log(4). The genome size for the calculation was obtained from the Animal Genome Size Database [Gregory, 2017], in which the reported genome sizes for the *Amphiprion* genus ranged from 792 to 1,200 Mb. The genome size of the *A. frenatus* individual was subsequently estimated from the genomic data, by dividing the number of error-free 18-mers by their peak coverage depth.

MP reads were processed at Fasteris SA (Geneva). Because MP libraries can have a relatively low total diversity, the dataset was screened for paired-reads sharing the exact same sequences on the first 30 bases of both sides. This can be expected due to PCR artifact and only one of the copies was kept to obtain unique pairs. The dataset was additionally screened to remove reads containing empty inserts. The linker sequences were searched and trimmed in the unique and not-empty pairs. Sickle was used to remove the remaining low-quality bases (parameters: --qual-threshold 25 --length-threshold 80). The quality of the resulting MP was verified with FASTQC (v0.11.2) [Andrews, 2010].

PacBio long reads were corrected with proovread (v2.12) [Hackl *et al.* 2014] using trimmed and error-corrected PE reads. This method allows to increase SMRT sequencing accuracy, which is substantially lower compared to Illumina technologies [Goodwin *et al.* 2016]. Because two different individuals were sequenced, proovread also corrects the possible polymorphism based on the Illumina-sequenced individual. This will remove possible errors due to the sequencing of different individuals for the genome assembly [Zhu *et al.* 2016].

### Nuclear genome assembly

Trimmed MP and PE reads resulting from the two strategies of read correction were assembled using both Platanus (v1.2.1) [Kajitani *et al.* 2014] and SOAPdenovo2 (v2.04.240) [Luo *et al.* 2012]. One of the advantages of Platanus is its automatic optimization of all parameters, including *k*-mer size. In SOAPdenovo2, assemblies were performed with a *k*-mer size of *k*=35 and *k*=63. The two values were chosen to span a large range, with the lower being comparable to the starting *k*-mer size of Platanus, and the larger being close to the best *k* proposed by KmerGenie [Chikhi and Medvedev 2013].

Scaffolding and gap-closing were performed within the Platanus or SOAPdenovo2 pipelines. For scaffolding, both short-insert and long-insert libraries were used. The best genome assembly was selected by investigating assembly statistics (N50, maximum scaffold length, number of scaffolds, gap number). The best genome assembly was reached using the reads corrected with ALLPATHS-LG modules and assembled with Platanus. Because of the substantial better quality of Platanus assemblies over the SOAPdenovo2 ones, we decided not to perform SOAPdenovo2 assemblies by progressively increasing *k-*mer sizes.

We further closed gaps in the resulting best assembly using the corrected PacBio long reads with PBJelly2 (v14.1) [English *et al.* 2012]. The script FakeQuals.py was used to set a quality score of 40 to each base in each scaffold. The mapping of long reads on the genome in PBJelly was performed by blasr [Chaisson *et al.* 2012], with the parameters set as following: -minPctIdentity 70 -SdpTupleSize 11 -nCandidates 20. The other parameters were left as default. Scaffolds smaller than 1 kb were removed from the final assembly.

### RNA extraction, library construction, sequencing and read processing

An additional individual of *A. frenatus* was obtained from a local aquarium shop for RNA sequencing to improve gene annotation of genome assembly [Denton *et al.* 2014]. RNA was extracted from liver with RNeasy Mini Kit (Qiagen, Hilden, Germany) following manufacturer’s instructions. The total amount of RNA in each sample and its quality was measured using Fragment Analyzer Automated CE System.

A strand-specific cDNA library was prepared using TruSeq Stranded mRNA Sample Prep Kit (Illumina) from an initial amount of total RNA of 1 μg, and following manufacturer’s instruction. The concentration, purity, and size of the library were tested using Fragment Analyzer Automated CE System. The library was sequenced on 1 lane of Illumina HiSeq2000 at the LGTF (run type: paired-end reads, read length of 100). Obtained PE reads were trimmed with Sickle, with the following parameters: --qual-threshold 20 --length-threshold 20.

### Nuclear genome validation

We investigated the quality of the assembled genome by evaluating the mapping rates of the PE and MP libraries using BWA (v0.7.12) [Li and Durbin, 2010], with default parameters. PE were subsampled and only the reads from a single Illumina lane were used. Prior to BWA mapping, MP were reversed to obtain the forward-reverse orientation with a homemade script. The RNA-Seq reads from *A. frenatus* were mapped to the genome with HiSat2 (v2.0.2, default parameters) [Kim *et al.* 2015]. Mapping statistics were summarized with BamTools stats (v2.3.0) [Barnett *et al.* 2011]. Insert sizes and read orientation were checked with Picard (v2.2.1, “CollectInsertSizeMetrics” tool) [http://picard.sourceforge.net].

The composition of the short scaffolds (< 1kb) removed from the final assembly was assessed using BLASTN (v2.3.30, https://blast.ncbi.nlm.nih.gov/Blast.cgi) against RefSeq database (Release 80, E-value cut-off of 10^−4^).

To further assess the quality of the assembly, a microsynteny analysis against *Oreochromis niloticus* genome (GCA_000188235.2) [Brawand *et al.* 2014] was performed with SynChro [Drillon *et al.* 2014]. We allowed for 5 to 10 intervening genes between gene pairs, as performed in Dibattista *et al.* [2016]. Finally, the completeness of the genome assembly was assessed with CEGMA (v2.3) [Parra *et al.* 2007].

### Nuclear genome annotation

Interspersed repeats and low complexity DNA sequences in the genome were identified with RepeatModeler (v1.08, engine ncbi) and soft-masked with RepeatMasker (v. 4.0.5) [Smit *et al.* 2013-2015]. *Ab initio* gene prediction was carried out with BRAKER1 [Hoff *et al.* 2015]. RNAseq data of *A. frenatus* previously mapped with HiSat2 (see Genome Validation) was used within BRAKER1 to improve *ab initio* gene prediction. RNA-Seq data was subsequently assembled into transcripts with Cufflink (v.2.2.1, default parameters) [Trapnell *et al.* 2010]. These transcripts, together with the *ab initio* gene predictions and the proteomes of *Danio rerio* (GCA_000002035.3), *Oreochromis niloticus* (GCA_000188235.2) and *Stegastes partitus* (GCA_000690725.1) were used to provide evidence for the inference of gene structures. The different evidences were aligned on each genome and synthesized into coherent gene models with MAKER2 (v2.31.8) [Holt and Yandell 2011]. The quality of the annotation was assessed by investigating the annotation edit distance (AED), which is calculated by MAKER2 [Holt and Yandell 2011].

The completeness of the resulting gene models was assessed by comparing the length of the predicted proteins with the *O. niloticus* proteins length. We performed BLASTP searches against *O. niloticus* proteome (total of 47,713 proteins, E-value cut-off of 10^−6^), We assumed that the best blast hit was orthologous and calculated the difference in protein length. We also calculated the “query” (*A. frenatus*) and “target” (*O. niloticus*) coverage, as defined in NCBI Genome (Supplementary Figure S1 in https://www.ncbi.nlm.nih.gov/genome/annotation_euk/Oreochromis_niloticus/102/).

Functional annotation was performed with BLASTP searches against the SwissProt database (subset: metazoans proteins, downloaded on June 2016, total of 104,439 proteins), with an E-value cut-off of 10^-6^. We also blasted (BLASTP, E-value cut-off of 10^−6^) *A. frenatus* proteins against RefSeq database (subset Actinopterygii sequences, downloaded on June 2016, total of 175,995 sequences), which is less accurate than SwissProt but more comprehensive. To provide further functional annotations, we used InterProScan (v5.16.55.0) [Jones *et al.* 2014] to predict protein domains based on homologies with the PFAM database (release 28, 16,230 families) [Finn *et al.* 2016]. Gene Ontologies (GO) were annotated to each predicted protein by retrieving the GO associated to its best SwissProt hit. Additionally, GO associated to protein domains were annotated in the InterProScan pipeline (option-goterms).

The completeness of the genome annotation was investigated with BUSCO (v.2, datasets: Metazoan and Actinopterygii, mode: proteins) [Simão *et al.* 2015]. Additionally, we calculated the “query” and “target” coverage for *A. frenatus* proteins and their SwissProt hits (Supplementary Figure S1). Query coverage and target coverage were compared to *O. niloticus, Maylandia zebra* and *D. rerio* coverages retrieved from NCBI (https://www.ncbi.nlm.nih.gov/genome/annotation_euk/Oreochromis_niloticus/102/). The *C. austriacus* proteome was downloaded from http://caus.reefgenomics.org on January 2017. As for *A. frenatus*, protein sequences of *C. austriacus* were blasted against SwissProt metazoan, and “query” and “target” coverages were calculated.

### Mitochondrial genome reconstruction and annotation

We reconstructed the entire mitochondrial genome from a random subsample of 20 millions of the PE reads filtered with ALLPATH-LG. To do so, we followed the baiting and iterative mapping approach implemented in MITObim (v1.9) [Hahn *et al.*, 2013]. We checked for the consistency of the outputs of two reconstruction methods. First, we used as reference a previously published complete mitochondrial genome of *A. frenatus* that we retrieved from GenBank (GB KJ833752) [Li *et al.*, 2015]. Alternatively, we also worked using a conspecific barcode sequence (*i.e.* COI gene) as a seed to initiate the process (GB FJ582759) [Steinke *et al.*, 2009]. The circularity of the sequence was manually inferred and the reads of the pool were mapped back onto the resulting mitochondrial genome to check for the reconstruction success and to assess coverage using Geneious (v10.2.2) [Kearse *et al.*, 2012]. We used MitoAnnotator and the MitoFish online database to annotate the inferred mitochondrial genome [Iwasaki *et al.*, 2013].

## Results and Discussion

### Nuclear genome sequencing and assembly

We obtained 534.9 million raw PE reads, corresponding to 108 Gb and 126X coverage. ALLPATH-LG error correction led to a total of around 105 Gb, while Quake strategy led to a smaller number of bases in total (76 Gb). This difference is due to the strict Phred score threshold that was set to 30 during the trimming, which caused the removal of most of the reads from the dataset. For MP data, we obtained 123.4 million raw pairs, which decreased to 57.1 million reads after filtering (9.6 ×). For PacBio long reads, we obtained 552,529 reads after correction with Illumina PE covering 1.8 Gb (2.2X). A summary of the sequencing results is provided in Table 1.

**Table 1.** *A. frenatus* DNA sequencing statistics for Illumina short reads and PacBio long reads.

| **Paired-end Library (350 bp insert)** |  |  |  |  |  |
| --- | --- | --- | --- | --- | --- |
|  | # Pairs | # Orphan | Mean Length | Total # bases | Coverage<sup>1</sup> |
| Raw data | 534 892 676 | 0 | 101 bp | 108 048 320 552 | 126.1 X |
| ALLPATH-LG | 507 172 071 | 22 941 362 | 101 bp | 104 765 835 904 | 122.2 X |
| QUAKE | 335 811 384 | 88 162 294 | 99.9 bp | 75 932 521 405 | 88.6 X |
| **Mate pairs library (3 kb insert)** |  |  |  |  |  |
|  | # Pairs | # Orphan | Mean Length | Total # bases | Coverage<sup>1</sup> |
| Raw Data | 123 437 124 | 0 | 125 bp | 30 859 281 000 | 36.0 X |
| Unique Reads | 96 476 505 | 0 | 125 bp | 24 119 126 250 | 28.1 X |
| Final reads | 57 140 128 | 3 245 451 | 70.3 bp | 8 260 918 652 | 9.6 X |
| **PacBio reads** |  |  |  |  |  |
|  | # Reads | Mean Length | Median Length | Total # bases | Coverage<sup>1</sup> |
| Raw Data | 660 691 | 3 480 bp | 2 751 bp | 2 299 043 897 | 2.7 X |
| Corrected reads | 552 529 | 3 634 bp | 3 009 bp | 1 898 588 929 | 2.2 X |

<sup>1</sup>: coverage is calculated with an estimate genome size of *A. frenatus* of 857 Mb.

The frequency of *k*-mers in ALLPATH-LG module estimated a genome size of 857 Mb, while in Quake strategy the estimated genome size was 820 Mb. The C-values for *A. frenatus* are not known, but available C-values for *A. perideraion* range from 0.81 (792 Mb) to 1.22 (1.2 Gb; from Animal Genome Size Database, Gregory, 2017).

We selected the best genome assembly by investigating assembly statistics (N50, maximum scaffold length, number of scaffolds, gap number). The best assembly was achieved with ALLPATH-LG corrected PE and assembled with Platanus (Supplementary Table S1). After further gap-closing with PacBio long reads, the final assembly included 17,801 scaffolds (> 1 kb), which covered a total length of 803.3 Mb (Table 2). Although the number of scaffolds is still important, 95% of the assembly is contained in less than 5000 scaffolds (Figure 1), and the N50 and N90 statistics are respectively 244.5 and 48.1 kb. The longest scaffold measures 1.7 Mb and the assembly contains 1.5% of gaps (Table 2). These values are comparable to other published bony fish genomes [Austin *et al*. 2015, Nakamura *et al*. 2013, DiBattista *et al.* 2016]. For example, the genome of the Pacific bluefin tuna (*Thunnus orientalis*) is composed of 16,802 scaffolds (> 2 kb), with a N50 of 137 kb and the longest scaffold of 1 Mb [Nakamura *et al.* 2013]. Similarly, the draft genome assembly of the blacktail butterflyfish (*Chaetodon austriacus*) is composed of 13,967 scaffolds (> 200 bp), with a N50 of 150.2 kb and 6.85% of gaps [Dibattista *et al.* 2016]

**Table 2.** Assembly statistics for *A. frenatus* assembly.

|  | Contigs | Scaffolds |
| --- | --- | --- |
| Total Assembly Size | 790 913 538 | 803 326 750 |
| # Sequences | 102 763 | 17 801 |
| Longest Sequence [bp] | 152 672 | 1 727 223 |
| Average length [bp] | 7 696 | 45 128 |
| GC content | 39.6 % | 39.6 % |
| non-ATGC characters [%] | 0 | 1.5 % |
| Number of gaps | 0 | 84 962 |
| Sequences >= 10 Kb | 25 615 | 5 646 |
| Sequences >= 1 Mb | 0 | 10 |
| N50 index (count) | 14 928 (15 134) | 244 530 (1 001) |
| N90 index (count) | 3 644 (55 579) | 48 151 (3 637) |

**Figure 1:**
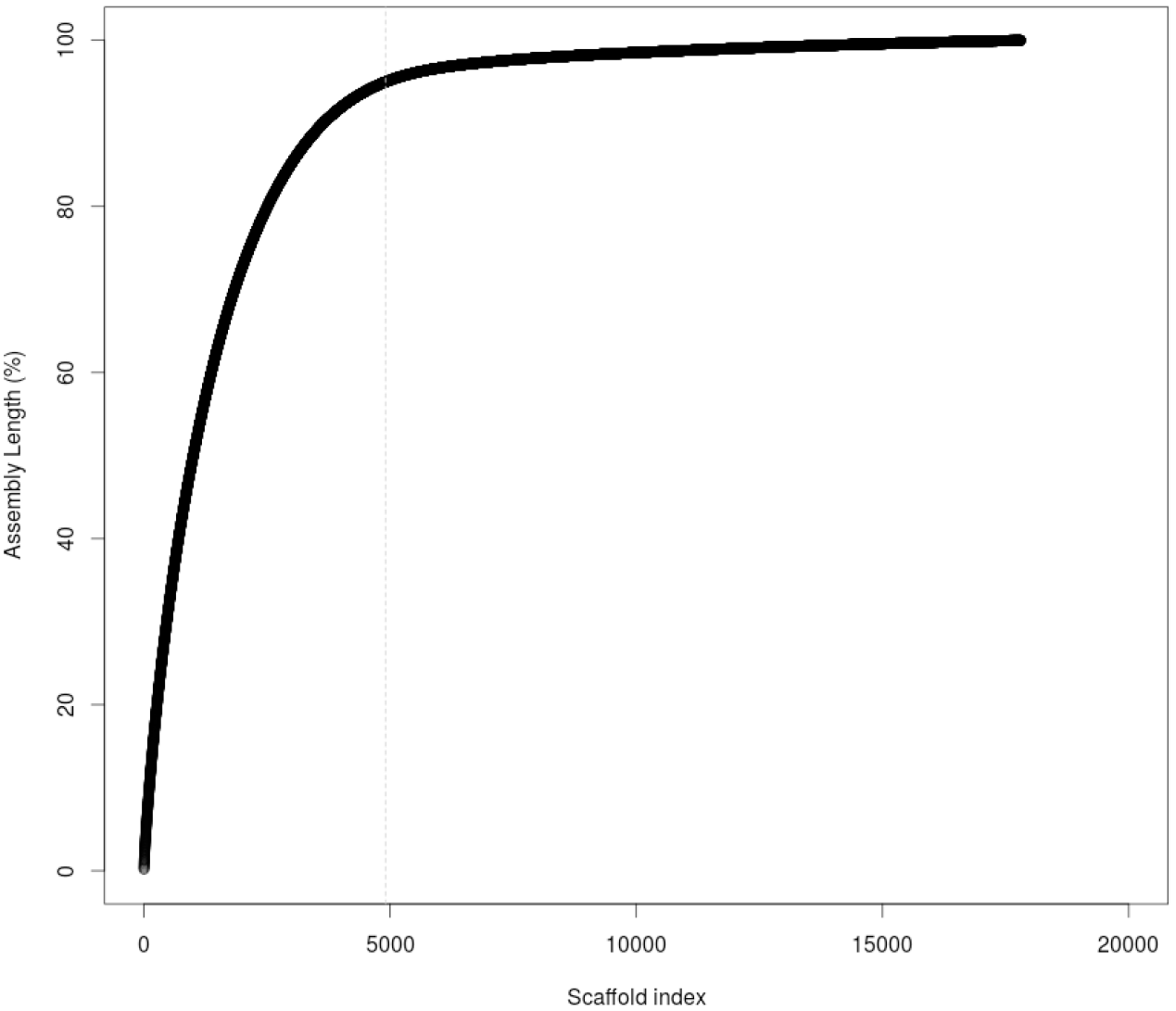
Cumulative length of *A. frenatus* assembly. Scaffolds are sorted from the longest to the shortest along the horizontal axis. The vertical dotted line indicates the number of scaffolds containing 95% of the assembly.

### Nuclear genome validation

The overall mapping rates for PE, MP and RNA-Seq PE data were of 99.4 %, 98.2 and 92.3 %, respectively (Table 3). The distribution of insert sizes estimated from the mapping was similar to the distribution obtained during the library preparation (Table 3 and supplementary figure S2). Some larger inserts were estimated for RNA-Seq data and are explained by the presence of introns in the genome. The high mapping rates and expected insert sizes reflect an overall good assembly. This is especially true for RNA-Seq data, which was obtained independently and was not used during genome assembly.

**Table 3.**
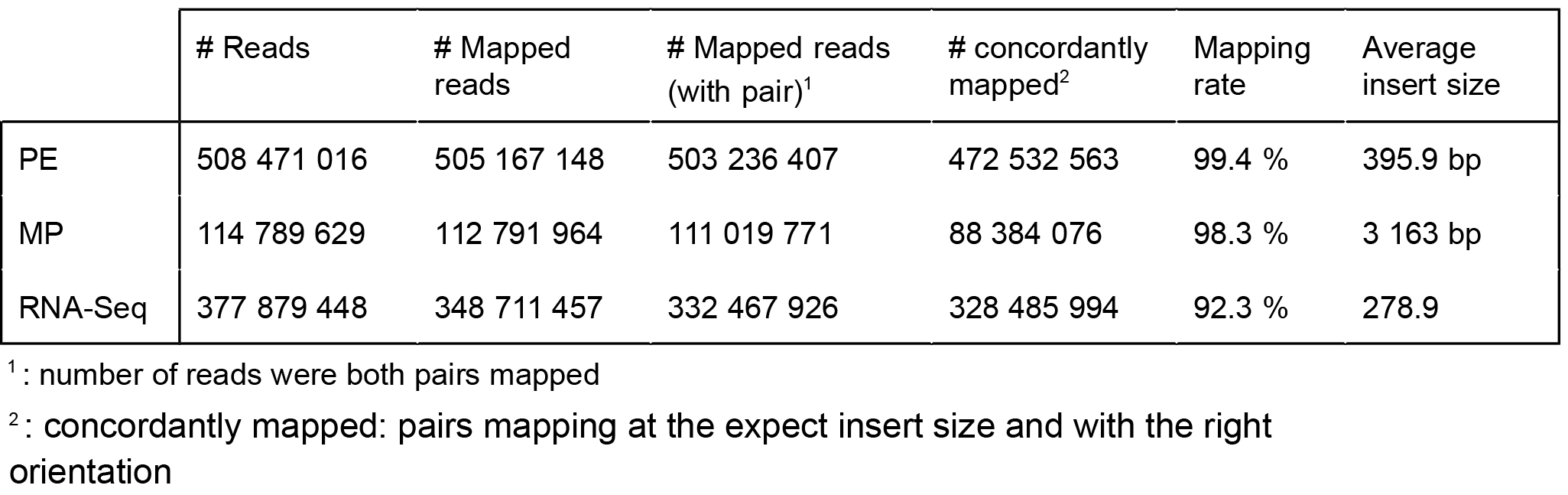
Mapping rates for paired-end (PE) reads, mate-pairs (MP) and RNA-Seq data. PE and MP reads were mapped with BWA. RNA-Seq data was mapped with HiSat.

**Table 4.** BUSCO results for completeness of *A. frenatus* genome assembly and annotation.

|  | Actinopterygii |  | Metazoan |  |
| --- | --- | --- | --- | --- |
|  | Counts | Percent | Counts | Percent |
| Complete, single-copy | 3542 | 77.3 % | 821 | 83.9 % |
| Complete, duplicated | 737 | 16.1 % | 104 | 10.6 % |
| Total Complete | 4279 | 93.3 % | 925 | 94.6 % |
| Fragmented | 150 | 3.3 % | 19 | 1.9 % |
| Missing | 155 | 3.4 % | 34 | 3.5 % |

We omitted around 1.5 million scaffolds of small size (< 1 kb; 21 % of the assembly) from the final assembly, the majority of which (89.5 %) had no matches in the RefSeq database.

We used CEGMA to assess the completeness of the assembly, which resulted in 99% of the core genes being either completely or partially represented in our assembly (Supplementary Table S2). The microsynteny analysis of *A. frenatus* and *O. niloticus* genome gave 2,383 syntenic blocks containing a total of 13,821 (5 intervening genes) and 13,847 (10 intervening genes) genes (Supplementary Table S3).

### Nuclear genome annotation

The amount of repetitive elements in our *A. frenatus* genome was 27.83 %. With a combined approach of *ab initio* gene prediction and evidence-based homology, we identified 26,917 genes coding for 31’054 predicted proteins (Supplementary Table S4). All the genes were predicted in a total of 6,497 scaffolds composing the 93% of the total assembly length. The quality of the models is satisfying, with an average and median annotation edit distance (AED) of 0.19 and 0.14 respectively (Supplementary Figure S3).

The lengths of *A. frenatus* predicted proteins were compared with the corresponding *O. niloticus* best hits. A total of 28,964 predicted proteins aligned with 22,110 *O. niloticus* targets and around half (56.3%) had less than 50 amino acids length differences with the target proteins (Supplementary Figure S4, left panel). Additionally, for 20,411 *A. frenatus* proteins, the “query” coverage was higher than 90%. Similarly, the “target” coverage was higher than 90% in 17,419 cases (Supplementary Figure S4, right panel).

The majority of the genes (86.5 %) returned a match to SwissProt metazoan proteins. This number further increased to 94.9% when we blasted our data against RefSeq database. Protein domain annotation was possible for 25,002 genes with 5,397 domains and 2,999 gene ontologies associated with these domains. A total of 17,788 gene ontologies were also mapped to 25,862 proteins (Supplementary Table S5).

The largest number of genes annotated with RefSeq is explained by a lower divergence between *A. frenatus* and the Actinopterygii species selected from the RefSeq database. Indeed, most of the best SwissProt database hits were obtained with human sequences (Supplementary Figure S5). This lower divergence also explains the higher identity for the matches obtained with RefSeq database (82.1% of average identity) compared with the SwissProt database (61.5% of average identity). Similarly, only 4,607 proteins had identity higher than 80% with proteins from SwissProt, while this number increased to 19,322 for RefSeq (Supplementary Figure S6). When comparing the completeness of *A. frenatus* gene models with other Actinopterygii species, we obtained results similar than *O. niloticus*, *M. zebra* and *D. rerio*, with 14,441 *A. frenatus* proteins having a “query” coverage larger than 90% and 12,605 proteins having a “target” coverage higher than 90%. Similar results were obtained for *C. austriacus* genome (Figure 2).

BUSCO analyses were performed to assess the completeness of *A. frenatus* assembly and annotation. For metazoan BUSCOs, 3.4 % of the genes were missing, while for Actinopterygii BUSCOS, 3.5 % of the genes were missing (Table 5).

**Figure 2.**
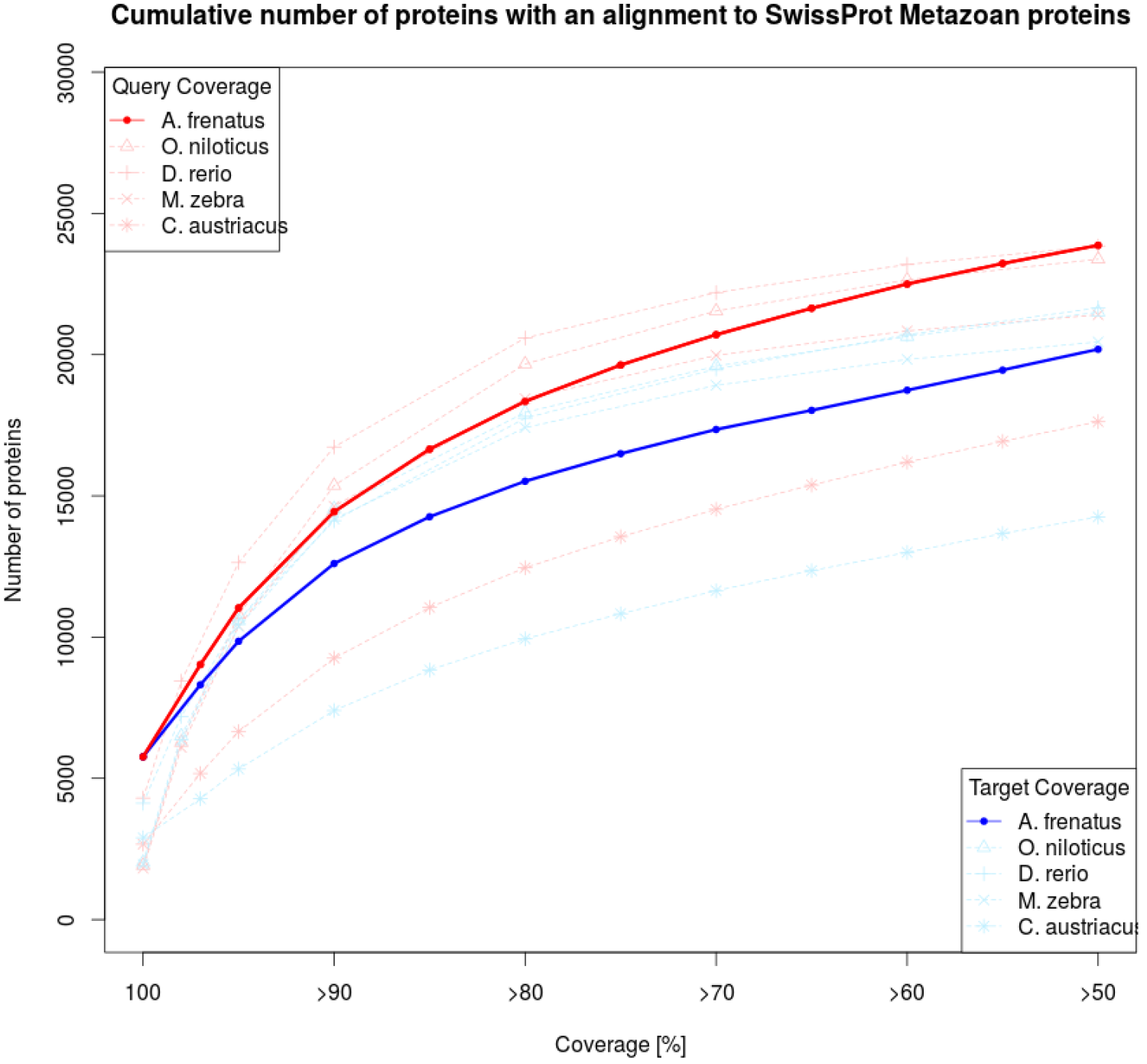
Query (red) and target (blue) coverage for *A. frenatus*, *O. niloticus*, *D. rerio*, *M. zebra* and *C. austriacus* proteins and their best SwissProt hit proteins.

### Mitochondrial genome reconstruction

We successfully reconstructed the complete mitochondrial genome of *A. frenatus*. The two methods used gave highly congruent results with each other. The mapping of the reads onto the inferred sequences led to a mean coverage of 20X (4X to 35X) and confirmed that the sequence could be unambiguously reconstructed. The inferred consensus sequence had a total length of 16,740 bp, which is slightly shorter than the 16,774 bp of the two available *A. frenatus* mitochondrial sequences (GB KJ833752, Li *et al.*, 2015 and GB LC089039, Thongtam na Ayudhaya *et al*., unpublished data). Its H-strand nucleotide composition is A: 29.6%, T: 25.7%, C: 29.3% and G: 15.4% and its GC content is 44.7%. This circular genome has a structure that is typical of fish mitochondrial genomes. It contains 13 protein-coding genes, 22 transfer RNA (tRNAs) genes, 2 ribosomal RNAs (rRNAs), 1 control region (D-loop) plus another 33 bp short non-coding region (OL) located between the tRNA-Asn and the tRNA-Cys (see Supplementary Table S6 and Supplementary Figure S7 for details). Pairwise differentiation between the three mitochondrial genomes available ranged from 0.77% to 2.0% suggesting an interesting amount of intra-specific variation in *A. frenatus*.

## Conclusion

Here we presented the first nuclear genomic resource for a clownfish species. Despite the fragmented nature of our assembly, the overall quality and completeness of the tomato clownfish nuclear genome are satisfying and comparable to other recent bony fish genome assemblies.

The genome that we present here, along with further sequencing of additional species and possible sequencing refinement, provides a new resource for future investigations of clownfishes adaptive radiation and their particular life history traits. It will also enable a deeper understanding of the origin of the mutualistic interactions with sea anemones by opening the way for comparative genomics analyses, which could allow the identification of the genomic bases of clownfishes adaptive radiation. Additionally, the resource that we present here will allow the design of new phylogenetic or population genomics markers that can be useful to study clownfish and damselfish evolution.

## Data accessibility

Raw Illumina and PacBio reads will be available in the Sequence Read Archive (SRA) in the NCBI database. The assembled nuclear genome, mitogenome, and their annotation will be available in DRYAD repository.

## Acknowledgments

We would like to thank the Vital-IT infrastructure from the Swiss Institute of Bioinformatics for the computational resources and to the Lausanne Genomic Technologies Facility for the sequencing.

